# DUETT quantitatively identifies known and novel events in nascent RNA structural dynamics from chemical probing data

**DOI:** 10.1101/458703

**Authors:** Albert Y. Xue, Angela M Yu, Julius B. Lucks, Neda Bagheri

## Abstract

**Motivation:** RNA molecules can undergo complex structural dynamics, especially during transcription, which influence their biological functions. Recently developed high-throughput chemical probing experiments study RNA cotranscriptional folding to generate nucleotide-resolution ‘reactivities’ for each length of a growing nascent RNA and reflect structural dynamics. However, the manual annotation and qualitative interpretation of reactivity across these large datasets can be nuanced, laborious, and difficult for new practitioners. We developed a quantitative and systematic approach to automatically detect RNA folding events from these datasets to reduce human bias/error, standardize event discovery, and generate hypotheses about RNA folding trajectories for further analysis and experimental validation.

**Results:** <u>D</u>etection of <u>U</u>nknown <u>E</u>vents with <u>T</u>unable <u>T</u>hresholds (DUETT) identifies RNA structural transitions in cotranscriptional RNA chemical probing datasets. DUETT employs a feedback control-inspired method and a linear regression approach and relies on interpretable and independently tunable parameter thresholds to match qualitative user expectations with quantitatively identified folding events. We validate the approach by identifying known RNA structural transitions within the cotranscriptional folding pathways of the *Escherichia coli* signal recognition particle (SRP) RNA and the *Bacillus cereus crcB* fluoride riboswitch. We identify previously overlooked features of these datasets such as heightened reactivity patterns in the SRP RNA about 12 nucleotide lengths before base pair rearrangement. We then apply a sensitivity analysis to identify tradeoffs when choosing parameter thresholds. Finally, we show that DUETT is tunable across a wide range of contexts, enabling flexible application to study broad classes of RNA folding mechanisms.

**Availability:** https://github.com/BagheriLab/DUETT

## 1 Introduction

RNA molecules play diverse functional roles ranging from catalysis of splicing and peptide bond formation, regulation of mRNA processing and gene expression, and molecular scaffolding and localization (Sharp, 2009; Cech and Steitz, 2014). These functions are in turn mediated by RNA structures that form in complex cellular environments. RNA structures are diverse and can prohibit or promote interactions with other RNAs, proteins, and metabolites to enable a broad range of RNA function. For example, bacterial RNA structures inhibit transcription elongation (Jagadeeswaran *et al.*, 2010), translation initiation (Afonin *et al.*, 2016), and RNA degradation (Hui *et al.*, 2015). In eukaryotes, there is growing evidence that RNA structure impacts gene expression processes (Rouskin *et al.*, 2014; Ding *et al.*, 2014; Spitale *et al.*, 2015; Talkish *et al.*, 2014). However, we know little about how newly synthesized, or nascent RNAs fold during transcription (Woodson, 2010; Pan and Sosnick, 2006). Due to the relative timescales of RNA folding and transcription, RNA molecules begin to fold as they emerge from RNA polymerase (Dethoff *et al.*, 2012) (Figure 1A). RNAs can transition between states in its cotranscriptional folding pathway that dictate RNA folding and function. For example, riboswitches dynamically alter their structure during transcription in response to ligand binding, leading to ligand-dependent structural, and regulatory switching (Smith *et al.*, 2009). In addition, there is emerging evidence that cotranscriptionally-formed RNA structures influence various processes in eukaryotes such as splicing (Shukla and Oberdoerffer, 2012) and 3’ end processing of histone mRNAs (Saldi *et al.*, 2016). There has been great interest in developing both computational and experimental approaches to uncover RNA cotranscriptional folding pathways and their implications for cellular RNA function.

**Figure 1.**
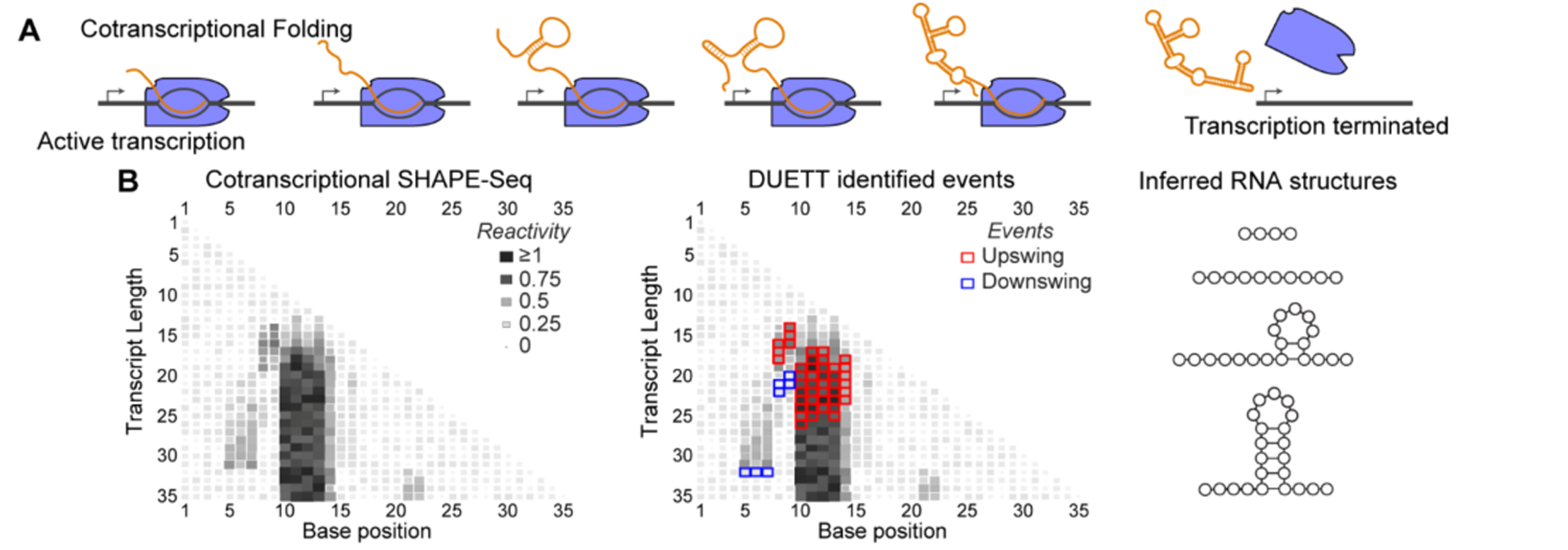
DUETT provides an automated and systematic method to detect cotranscriptional RNA folding events from SHAPE-Seq data. A) RNA can dynamically alter structure during transcription that affect downstream biological functions. B) Cotranscriptional SHAPE-Seq probes RNA structural properties by measuring reactivity patterns where high and low reactivity correspond to unstructured and constrained regions of RNA, respectively. DUETT is a flexible method to identify large reactivity changes that are indicative of RNA structural transitions. Here DUETT is applied to a mock dataset to identify an increase in reactivities in several consecutive nucleotides which is consistent with the typical ‘low-high-low’ pattern observed for RNA hairpin structures.

Recently developed experimental techniques can characterize cotranscriptional RNA folding at nucleotide resolution (Strobel *et al.*, 2017; Watters, Strobel, *et al.*, 2016) by utilizing high-throughput chemical probing of RNA structure (Strobel *et al.*, 2018). SHAPE (selective 2’-hydroxyl acylation analyzed with primer extension) are chemical probes that form adducts at the 2’-OH of each nucleotide (Merino *et al.*, 2005). When coupled with high-throughput sequencing, SHAPE experiments reveal detailed reactivity patterns that uncover RNA structural properties – highly reactive positions indicate lack of structure and lowly reactive positions indicate constraint due to structure or interaction with other binding partners (Bindewald *et al.*, 2011; Steen *et al.*, 2010; Watters, Yu, *et al.*, 2016). An experimental variant called cotranscriptional SHAPE-Seq was developed to probe the structure of each intermediate length RNA during transcription (Figure 1) (Strobel *et al.*, 2017; Watters, Strobel, *et al.*, 2016). This experiment results in a matrix of reactivities, where rows correspond to reactivity at each length of a growing nascent RNA chain, and columns represent reactivity changes at specific nucleotides across different RNA lengths (Figure 1B). Both dimensions of the reactivity matrix reflect possible changes in RNA structural state that can occur during transcription. For example, a decrease (or increase) in reactivity down a column highlights a possible folding (or unfolding) event during transcription.

However, analysis of the cotranscriptional reactivity matrices has been mostly qualitative, relying on manual identification of reactivity trends to identify key regions that have biological significance. As the number and complexity of these datasets grow, quantitative and automated techniques are needed to robustly identify patterns. This automated quantitative approach is challenging, as cotranscriptional SHAPE-Seq datasets with complete annotations and validated structures are not bountiful. This scarcity of “ground truth” examples presents difficulties when defining statistical models (Hogenboom et al., 2011) and prohibits application of machine learning models (Margineantu et al., 2010), which have been implemented in other applications such as emulating visual detection of meaningful reactivities as labeled by experimental experts (Woods and Laederach, 2017). These limitations suggest that we require a systematic method to identify signatures of RNA structural dynamics from cotranscriptional reactivity datasets that use interpretable, user-guided rules.

To overcome this challenge, we sought to develop a quantitative and automated approach to identify trends in cotranscriptional SHAPE-Seq datasets. We designed a systematic detection method that remains user-tunable with an interpretable set of parameters to easily match qualitative user expectations with quantitatively identified folding events. Other computational approaches have successfully emulated subjective human-driven analyses such as extracting RNA design rules from a crowd-sourced game (Lee *et al.*, 2015). Due to the complexity of RNA structures and the flexibility of SHAPE-Seq applications/implementations, we opted to detect generic events. This philosophy is common in domains with poorly defined events, such as identifying unknown genomic deletions and insertions (Ye et al., 2009; Jiang et al., 2015).

We present a framework for detecting events in cotranscriptional SHAPE-Seq datasets termed <u>D</u>etection of <u>U</u>nknown <u>E</u>vents with <u>T</u>unable <u>T</u>hresholds (DUETT). DUETT detects swing events using a strategy inspired by proportional-integral feedback control (Jr. and Parker, 2007) and detects ramp events using linear regression. Swing events represent rapid reactivity changes that occur over a small number of transcript lengths. In contrast, ramp events represent slower changes that span many transcript lengths. DUETT provides automated threshold parameter optimization, but DUETT also allows user-defined parameter tuning to match a wide range of experimental contexts. We first define these methods and identify parameters that robustly identify known folding events within the cotranscriptional folding pathway of the *E. coli* Signal Recognition Particle (SRP) RNA. We extend the methodology to analyze the folding pathways of the *B. cereus* crcB fluoride riboswitch and corroborate previous manually-identified transitions. In both datasets, our analysis reveals unexpected behavior, such as subtle reactivity increases that consistently occur roughly 12 nucleotide lengths before a reactivity decrease, suggesting a highly-reactive transient state. Finally, we conduct parameter sensitivity analysis to explore the relationship between DUETT’s parameter values and detected events. The flexibility and interpretability of our approach enables the broad application of DUETT to many high-throughput experimental systems that require event detection.

## 2 Methods

### 2.1 Event detection

Structural events are characterized by significant reactivity changes at specific nucleotide positions, across sequential transcript lengths. We consider two common, yet distinct, phenomena in cotranscriptional SHAPE-Seq datasets: swing and ramp events. These two qualitatively different classes of events motivate separate detection methods for each event type. DUETT methods and motivations are located in Supplementary Materials. Assumptions are explicitly listed alongside their design consequences in Supplementary Table S1.

### 2.2 Application to cotranscriptional SHAPE-Seq datasets

We applied DUETT to two RNA sequences characterized by previous cotranscriptional SHAPE-Seq experiments (Watters, Strobel, et al., 2016): the E. coli. SRP RNA and the B. cereus crcB fluoride riboswitch. These published datasets were obtained from the Small Read Archive (SRA) (http://www.ncbi.nlm.nih.gov/sra) with the BioProject accession code PRJNA342175. The data was processed with Spats v1.01 (https://github.com/LucksLab/spats/releases/) and the reactivity calculation scripts are located at https://github.com/LucksLab/Cotrans_SHAPESeq_Tools/releases/.

## 3 Results

We validated DUETT by identifying known cotranscriptional structural events that occur in the folding pathways of two RNA molecules: the *E. coli*. signal recognition particle (SRP) RNA and the *B. cereus*. fluoride riboswitch (Watters, Strobel, et al., 2016; Batey et al., 2000; Wong *et al.*, 2007). We used the automated approach to select *PIR* threshold parameters and manually selected the same linear ramp thresholds across all datasets. During the automated search, the increasing *PIR* thresholds cause the number of detected events to rapidly decrease until reaching an elbow (Supplementary Figure S2). The point closest to the origin lies near the elbow and corresponds to the optimized threshold values, similar to how numbers of clusters are chosen in clustering algorithms (Gao *et al.*, 2018). We applied DUETT on each of three experimental replicates for each system and retained events conserved across all replicates to decrease the likelihood of spurious events. This approach creates similar results as averaging all replicate datasets before applying DUETT (Supplementary Figure S3) but avoids scenarios where a single replicate has anomalous values. DUETT identified both known and novel structural events, and we propose novel hypotheses for further study. We discover patterns and events that are difficult for a human to identify. We conclude with parametric sensitivity analysis to explore the relationship between user-defined threshold parameters and observed events.

### 3.1 Validation on *E. coli* SRP RNA cotranscriptional SHAPE-Seq datasets and identification of previously unidentified reactivity patterns

Previous studies have shown that during transcription, the E.coli SRP RNA forms a transient 5’ hairpin (H1) that rearranges into a long helical structure with a hairpin loop and multiple inner loops (Batey *et al.*, 2000; Wong *et al.*, 2007; Watters, Strobel, *et al.*, 2016), which we label H2-H5 (Figure 2). Several of these transitions have been validated by prior bulk studies (Wong *et al.*, 2007; Watters, Strobel, *et al.*, 2016), and by single molecule optical trapping experiments (Fukuda *et al.*, 2018). A previous analysis of cotranscriptional SHAPE-Seq datasets (Watters, Strobel, *et al.*, 2016) focused on manual annotation of patterns within the reactivity matrix. We thus sought to apply DUETT to these datasets to automatically identify reactivity changes that are reflective of these structural transitions.

Within the cotranscriptional SHAPE-Seq datasets, many upramps (positive-slope ramps) begin near the 3’ end of the nascent RNA shortly after the nucleotide’s (nt) transcription by RNAP (Figure 2). This close association suggests SHAPE adduct formation occurs almost immediately after exiting RNAP. Due to experimental limitations (Strobel *et al.*, 2018), these short RNA fragments are difficult to detect, leading to reduced reactivity signal. However, as the RNA elongates, SHAPE adducts at these same positions become increasingly detectable, which could lead to the presence of gradually increasing reactivity upward ramps in these regions. As a result, we infer that upramps close to the nt’s exit from RNAP are likely experimental artifacts due to their position near the 3’ end of the RNA.

**Figure 2.**
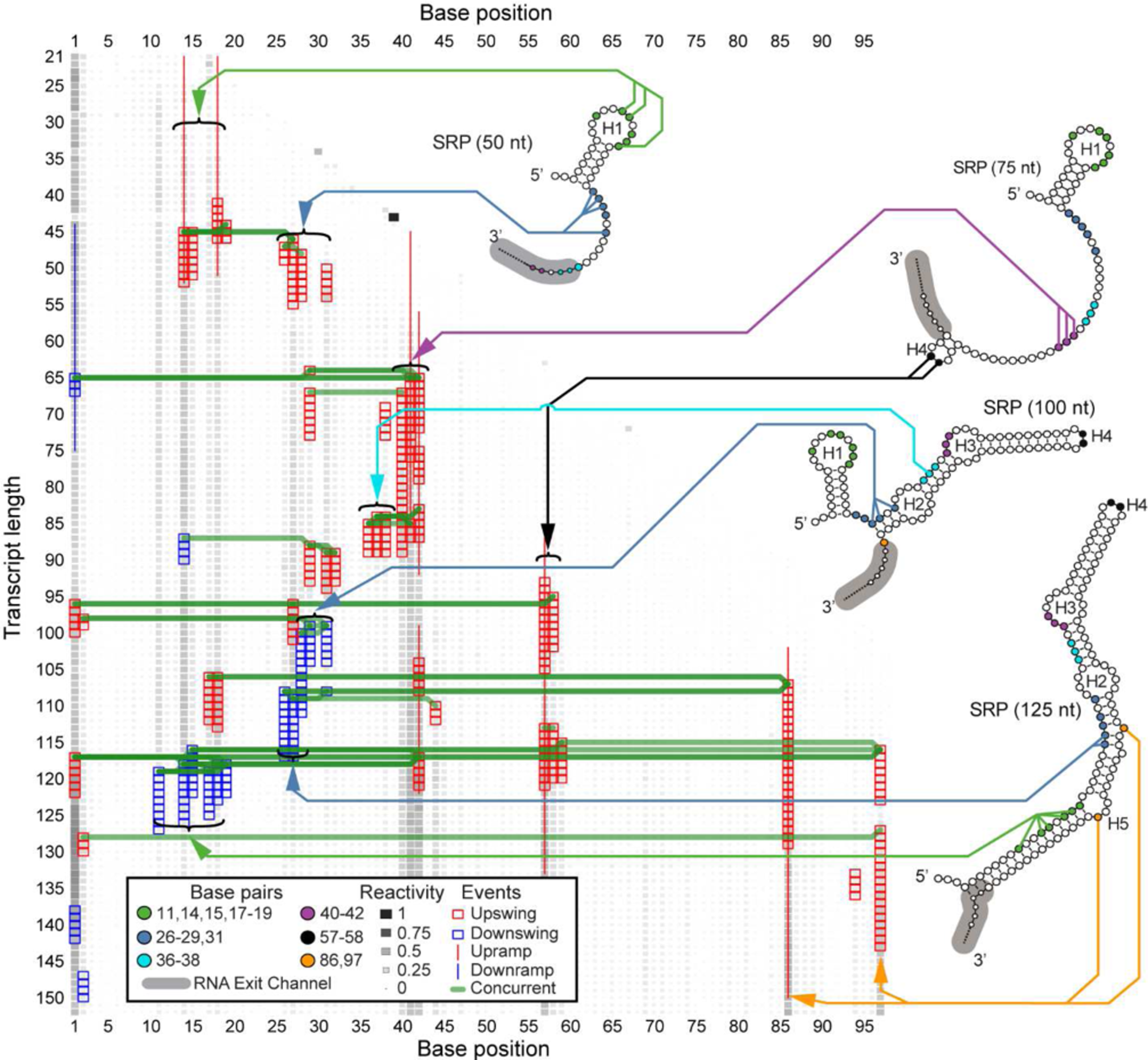
DUETT identifies known RNA folding events in the cotranscriptional SHAPE-Seq reactivity matrix derived from the *E. coli* SRP RNA. Four previously proposed intermediate structural conformations of SRP RNA are shown with arrows linking specific bases to identified reactivity changes. DUETT identifies multiple instances of hairpin formation/rearrangement and we propose structural explanations for previously unidentified novel events. DUETT displays detected swing and ramp events as a colored box and line, respectively, with red and blue denoting an increase and decrease, respectively. A green line connects concurrent events between different nucleotides. The RNA polymerase exit channel footprint that occludes the last 5 RNA nucleotides is annotated in grey. Nucleotides that participate in identified SHAPE-Seq reactivity transitions are color coded. RNA structures are from Figure 2 of (Watters, Strobel, *et al.*, 2016) with helices H1-H5 annotated. SHAPE reactivity is normalized to lie in between the range 0-1 and shown in greyscale and box area.

#### 3.1.1 DUETT identifies expected H1 formation and rearrangement

The major cotranscriptional rearrangement event for the *E. coli* SRP RNA occurs when H1 refolds into the final extended structure (Yu *et al.*, 2018; Fukuda *et al.*, 2018). Corresponding to the formation of H1, DUETT identified upswings in bases 14-15 and 18-19 around nascent RNA length 45 nt, and upramps that conclude around length 50 nt (Figure 2). These positions remain unpaired in the intermediate H1 hairpin, validating the correspondence between upswings/upramps and the formation of unpaired, reactive regions. This hairpin rearranges into the long helical structure, which DUETT identifies as downswing events between lengths 116 and 127 nt in bases 11, 14-15, and 17-19. These identifications are consistent with recent computational modeling (Yu *et al.*, 2018) and single molecule optical tweezer experiments (Fukuda *et al.*, 2018) that propose the rearrangement of H1 to occur in the window that DUETT detects.

#### 3.1.2 Multiple expected reactivity changes further validate DUETT

Another key feature of the *E. coli* SRP RNA cotranscriptional folding pathway is the formation of native base pairs and loops after the formation of H1, but before the rearrangement of H1 into the final native structure (Wong *et al.*, 2007; Watters, Strobel, *et al.*, 2016; Fukuda *et al.*, 2018; Yu *et al.*, 2018). DUETT identifies expected reactivity signatures of bases 26-29 and 31 through length 100 nt. Upon initial transcription, bases 26-28 and 31 have upswings corresponding to their unpaired state, and bases 28-29 and 31 have downswings at 100 nt that agree with the previously proposed 100 nt structure in which these nucleotides are paired (Watters, Strobel, *et al.*, 2016). Identification of other unpaired positions provides additional validation of DUETT event detection. A cluster of upramps/upswings in bases 40-42 between lengths 55 and 90 nts corresponds to the unpaired, interior loop region in H3. Finally, bases 57-58, 86 and 97 have upramps/upswings immediately after transcription that corroborates their unpaired status as the apex nucleotides in the hairpin loop of the final rearranged structure, or within internal loops and bulges, respectively.

#### 3.1.3 Unexpected events highlight overlooked structural dynamics

While DUETT identified previously validated and observed reactivity changes within the SRP RNA cotranscriptional folding pathway, it also identified novel and unexpected events such as a downswing in base 14 and upswings in bases 36-38 at lengths 87 and 84 nts, respectively. These events are discordant with the previously proposed folding model of the SRP RNA: base 14 remains unpaired in H1 and bases 36-38 pair with bases 74-76 between lengths 75-100 nts. However, these detected events do not appear spurious. The downswing in base 14 is concurrent with other undetected downswings in neighboring bases 11 and 15 (Supplementary File 1), and the upswings in bases 36-38 are concurrent and qualitatively similar with one another. These unexpected events occur in the transition that forms the H2 and H3 loops and suggests a transient state that causes decreased reactivity in base 14 and increased reactivity in bases 36-38. In addition, base 40 was reported to be paired by length 100 nt (Watters, Strobel, *et al.*, 2016) that corresponds to an undetected downswing at 94 nt (Supplementary File 1). Though base 40 has lower reactivity than bases 41-42 (Supplementary File 1), its reactivity is higher than expected from a base pair. Our DUETT results suggest that U40 is more labile than previously reported (Watters, Strobel, *et al.*, 2016). We attribute this accessibility to U40’s position at a helix end as well as part of a GU pair with G72 in the native structure and these features are known to be generally less stable (Jaeger *et al.*, 1989; Papanicolaou *et al.*, 1984).

Another unexpected set of events include the downswings of bases 26-27 at length 108 nt, which is earlier than the stable rearrangement of H1 into the final helical structure. These downswings in bases 26-27 are concurrent with two upswing events at bases 44 and 86 and an upramp in base 86. Base 44’s upswing is likely a spurious event due to a replicate anomaly (Supplementary File 1) but base 86 is a bulged nt in the native structure that forms opposite of base pairs involving positions 27-28. This concurrency suggests that the bulge formation in base 86 occurs simultaneously with bases 26-27 pairing up and agrees with the previously proposed model (Watters, Strobel, *et al.*, 2016).

We also observed unforeseen upswings about 12 nt lengths before downswings, suggesting a highly-reactive transient state. Bases 17-18, 27, 29, and 31 exhibit upswings roughly 12 nt lengths before their downswing events (Figure 2). The order in which these downswings occurs is consistent with the order of transcription, suggesting an order of folding events based on initial exposure and transcription. Coincidentally, when bases 26-27 or bases 29-31 undergo stabilizing structural dynamics, the preceding upswing pattern occurs in the 3’ base(s) of that group. Similarly, the preceding upswing followed by downswing behavior in bases 17-18 and suggests that increased flexibility in the 3’ side transiently occurs before becoming stabilized. We note the difficulty in manually detecting these patterns of events, justifying our automated and systematic approach. These observations lead us to believe that the detected events are not spurious, lack an explanation by previously published studies, and highlight the discoveries enabled by our systematic method.

### 3.2 DUETT identifies known and novel structural transitions in the cotranscriptional folding pathway of a fluoride riboswitch

We next sought to determine if DUETT could identify events in the *B. cereus* fluoride riboswitch cotranscriptional SHAPE-Seq data (Watters, Strobel, *et al.*, 2016). The *B. cereus* fluoride riboswitch is an RNA sequence that lies in the 5’ untranslated region of the crcB gene and cotranscriptionally folds into mutually exclusive structures that regulate downstream transcription depending upon whether the fluoride anion is bound (Baker *et al.*, 2012). Previous cotranscriptional SHAPE-Seq experiments were done in either fluoride-positive (10 mM NaF) or fluoride-negative (0 mM NaF) conditions. Manual analysis revealed distinct reactivity patterns that are reflective of ligand binding, and the bifurcation of the folding pathway in a fluoride-dependent manner. We thus applied DUETT to both conditions to identify both known and potentially novel RNA folding events.

#### 3.2.1 Similarity of events before length 69 nt between conditions

Before the structural divergence at the nascent RNA length of 69 nt, DUETT-detected events agree with the proposed model that RNA structures are similar in both fluoride conditions (Watters, Strobel, *et al.*, 2016). Upswings in both conditions occur between 40 nt and 55 nt in bases 15-16, 24, 27, 30, and 33-34 (Figures 3 and 4), which confirm their unpaired state. Additionally, in the fluoride-positive condition, bases 12-13 and 16 have downswings around length 60 that suggest they form a critical pseudoknot, in which two helices are now interleaved, prior to the previously proposed length 69 nt structure that contains this pseudoknot. Only base 12 has a downswing around length 60 in the fluoride-negative condition, which could indicate a less stable pseudoknot and thus a less stable aptamer. Base 30 has a consistent upswing in both conditions between lengths 49 and 53 nt, and the previously proposed model (Watters, Strobel, *et al.*, 2016) suggests that bases 28-30 are paired within a hairpin stem spanning nts 28-37 before length 58 nt. This base 30 upswing occurs when the hairpin stem theoretically forms (lengths 51-54) and may indicate delayed hairpin stability due to its short 3 bps length. The detected events before length 69 nt reflect similar RNA structures that form independent of the presence of fluoride (Watters, Strobel, *et al.*, 2016).

**Figure 3.**
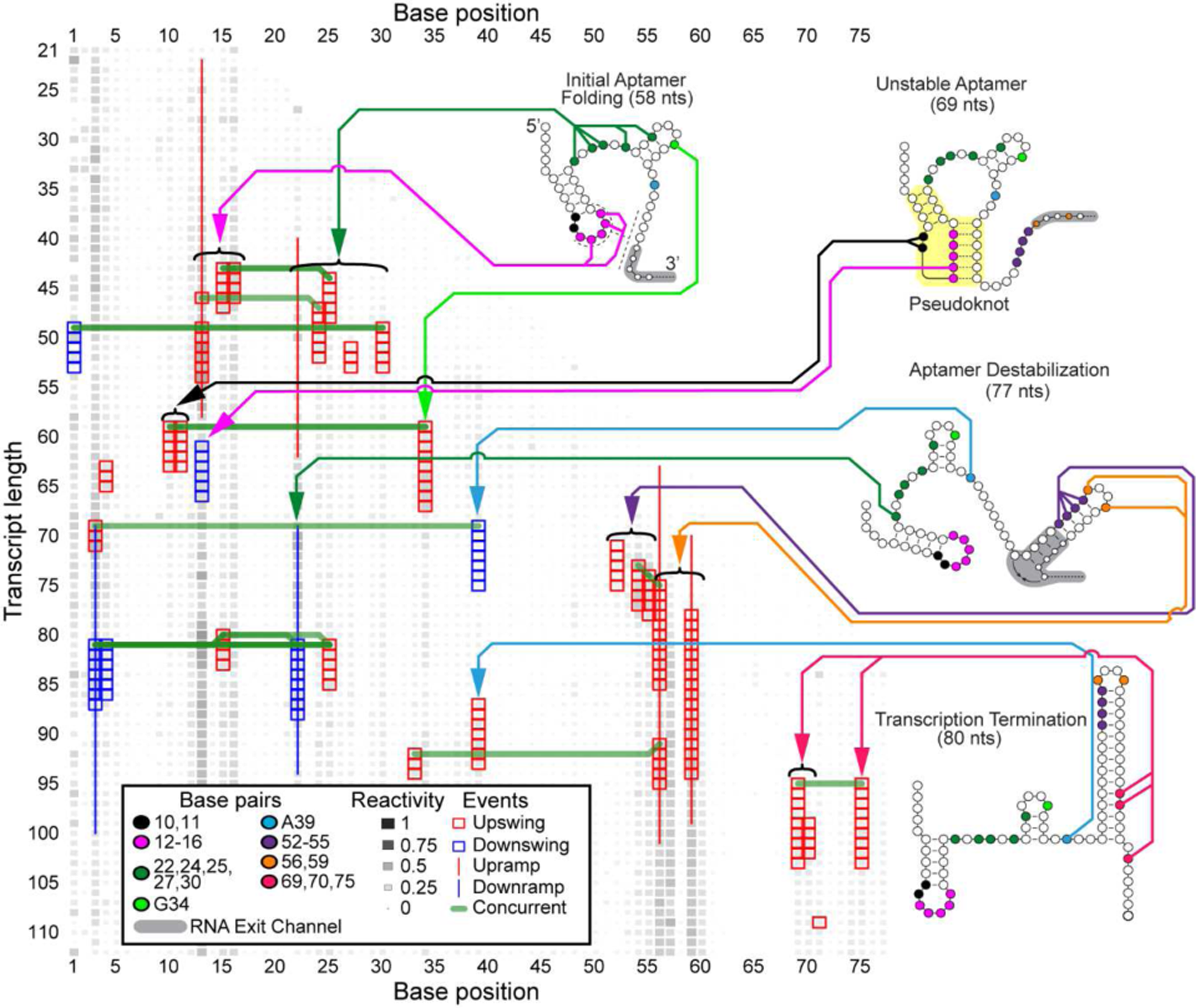
DUETT identifies changes in the *B. cereus* fluoride riboswitch without fluoride cotranscriptional SHAPE-Seq reactivity matrices. When comparing to events identified with the fluoride added condition (Figure 4), DUETT identifies multiple known and novel reactivity events, indicated by arrows to nucleotides participating in these events. The figure is annotated as in Figure 2. RNA structures and pseudoknot (highlighted in yellow) are drawn from Figure 6 of (Watters, Strobel, *et al.*, 2016).

**Figure 4.**
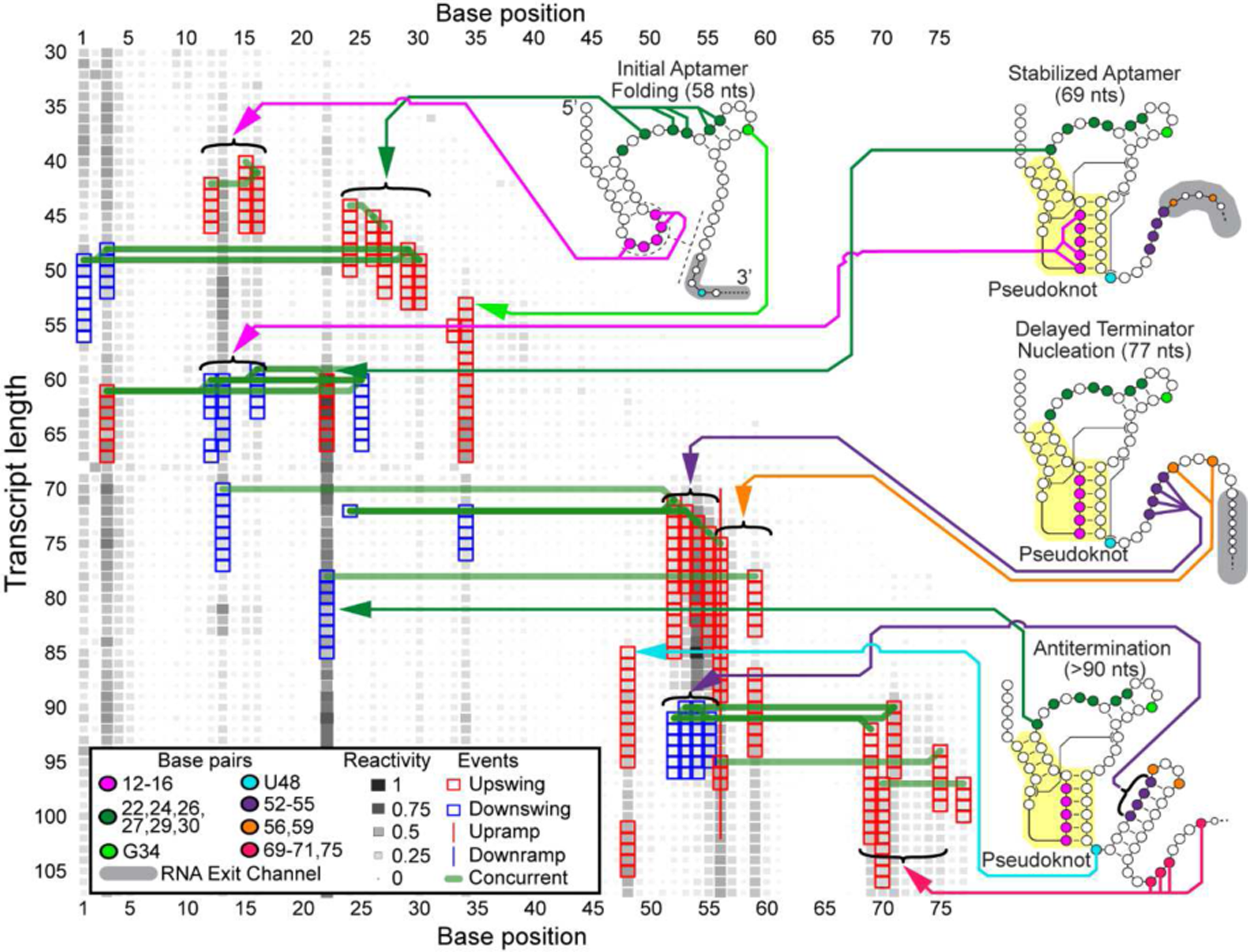
DUETT identifies changes in the *B. cereus* fluoride riboswitch with fluoride cotranscriptional SHAPE-Seq reactivity matrices. These results are compared to Figure 3 to identify structural divergences between the fluoride conditions. The figure is annotated as in Figure 3.

### 3.2.2 Identification of delayed terminator nucleation agrees with the previous riboswitch folding model

A key feature of the *B. cereus* crcB fluoride riboswitch folding pathway uncovered in previous work is a delay in the folding of the terminator RNA structure when fluoride is present (Watters, Strobel, *et al.*, 2016). Correspondingly, DUETT identified events that agree with this delayed terminator nucleation. Exclusively in the fluoride-negative condition, bases 12-16 were previously proposed to unpair by length 77 nt, allowing bases 52-55 to pair with bases 60-63 (Watters, Strobel, *et al.*, 2016). Both conditions exhibit upswings for bases 52-55 around lengths 70-75 nt due to different reasons: increased reactivity prior to hairpin formation in the fluoride-negative condition (similar to upswings observed before downswings in the SRP RNA dataset) and unpaired bases in the fluoride-positive condition. The fluoride-negative bases immediately decrease in reactivity (upon pairing off) while the fluoride-positive bases retain high reactivity (Supplementary Files 2 and 3). However, DUETT did not identify these downswings in the fluoride-negative case due to averaging reactivities within the sliding window that causes the upswing data to dominate identification for several following lengths; we further explore the effects of window size in Supplementary Materials. The delayed terminator nucleation in the fluoride-positive condition manifests as a series of downswings around length 90 nt exclusively in the fluoride-positive condition, which corresponds to forming the hairpin stem, but only after a delay of about 10 nts transcribed(Watters, Strobel, *et al.*, 2016). In addition, bases 56 and 59 exhibit upswings/upramps in both conditions, corroborating their unpaired nature within the loop of the terminator hairpin.

DUETT identified expected events that occur after terminator formation, when transcription is expected to halt exclusively in the fluoride-negative condition (Watters, Strobel, *et al.*, 2016; Ren *et al.*, 2012). As expected by their reactive nature in the fluoride-positive condition (Figure 4), bases 69-71 and 75 remain unpaired and contain upswings shortly after transcription. These upswings are expectedly missing in the fluoride-negative condition except for bases 69 and 70 (Figure 3), which exhibit unexpected upswing at length 95 and 99 nt, respectively. These events may have occurred because the RNAP transcribed past the expected termination site in the bulk cotranscriptional SHAPE-Seq experiment.

It was previously shown that the double mutant G69A, A70U prevents formation of the terminator stem, meaning that their pairing with bases 46-45 is a requisite of termination (Baker *et al.*, 2012; Watters, Strobel, *et al.*, 2016). A separate study, using a similar riboswitch sequence, found that a single long range reverse Hoogsteen base pair in the region shared by the aptamer and terminator stem area is pivotal in functional switching between termination and antitermination (Zhao *et al.*, 2017). These findings, coupled with the base 69 and 70 upswing, suggest that a subpopulation of fluoride riboswitch do not form the base pairs leading to increased reactivity and lost terminator function. However, the mechanism is unclear and requires further study. This previously overlooked observation demonstrates DUETT’s ability to identify interesting events for follow-up analysis.

### 3.2.3 Detected events in bases 10 and 48 confirm long-range interactions

DUETT corroborates two previously reported long-range interactions in the fluoride-positive condition: A10-U38 and A40-U48 (Watters, Strobel, *et al.*, 2016). These interactions were hypothesized to increase stability of the aptamer and persist only when fluoride binds (Watters, Strobel, *et al.*, 2016). In the fluoride-negative condition, we observe an upswing in both base 10 and base 11 at length 60 nt, which corresponds to the opening of the initial hairpin which could enable pseudoknot formation during aptamer formation (Figure 3). Conversely these upswings are absent in the fluoride-positive condition because the A10-U38 interaction prohibits increased SHAPE reactivity (Figure 4). The other long-range interaction, A40-U48, was proposed to unpair between the lengths 77 and 88 nt upon terminator hairpin nucleation (Watters, Strobel, *et al.*, 2016), which we also observe with an upswing in U48 at length 85 nt in the fluoride-positive data (Figure 4).

Additionally, A39 is situated between these two long-range interactions and exhibits an unexpected downswing and upswing in the fluoride-negative condition at 69 and 87 nt, respectively. After 77 nt, A39 exhibits structural divergence with an unexpected upswing at 87 nt. Previous NMR characterization of this system showed that A39 (A35 in their numbering) undergoes local structural dynamics when no fluoride is bound and is stabilized when fluoride is bound (Zhao *et al.*, 2017). These swing events may reflect those local structural changes. Conversely, the fluoride-positive condition lacks this upswing most likely due to the neighboring A10 U38 long-range interaction and continued aptamer presence. Altogether, DUETT detects swing events in bases 10-11, A39, and U48 that are consistent with the proposed aptamer stabilization via long-range interactions.

#### 3.2.4 DUETT identifies new A22 dynamics

DUETT identified SHAPE hyperreactivity in A22 via an upswing at length 60 nt in the fluoride-positive condition, which was associated with aptamer stabilization due to ligand binding (Watters, Strobel, *et al.*, 2016). This upswing is followed by a sharp downswing at length 78 nt, while another upswing shortly afterwards was undetected. The second upswing went undetected due to the short duration of the previous downswing, which causes high and low reactivity positions to lump together during the sliding window averaging. Afterwards, the reactivity plateaus at a high value comparable to length 69 nt. The fluoride-negative condition has similar dynamics but are less extreme and were detected as ramps (Supplementary Files 2 and 3) demonstrating that swing and ramp events differentiate small from large changes as intended. We conclude that base 22 has similar dynamics (except in magnitude) across both conditions until about length 90 nt where only the fluoride-positive condition exhibits the rebound upswing. The difference in magnitude of A22 reactivity between the conditions was previously concluded to be indicative of ligand-mediated aptamer stabilization and destabilization, respectively (Watters, Strobel, *et al.*, 2016). While the fluoride-negative downswing was thought to be due to aptamer destabilization, the analogous fluoride-positive downswing disagrees and lacks a mechanistic explanation since the aptamer is expected to be stable. A22’s complex behavior was overlooked earlier due to the visual upper limit (set as a reactivity value of 4) set in the original reactivity matrix figures (Watters, Strobel, *et al.*, 2016). While the upper limit simplifies data visualization, DUETT accounts for all magnitudes and partially insulated from disadvantages in human visualizations. Altogether, DUETT identified several expected structural differences between the fluoride conditions, and we generate multiple hypotheses to explain unknown or unexpected events.

### 3.3 Threshold parameters confer a tradeoff between true positive and false positive/negative events

DUETT balances the detection of true positive events with detection of erroneous false positive/negative events. Determining this balance highlights user preferences; if identifying small magnitude events is prioritized, then thresholds can be relaxed to increase finding true positive events at the cost of also increasing the number of false positives.

Parametric sensitivity analysis explores the true positive-false positive tradeoff. We demonstrate that large events are retained despite drastic threshold parameter adjustments. We highlight two scenarios in the *E. coli* SRP RNA dataset using a stringent and a lenient set of *PIR* thresholds (additional sensitivity tests are found in Supplementary Materials). The stringent *PIR* scenario (100% increase in each threshold) yielded fewer overall events (Supplementary Figure S4, right) relative to the original baseline (center). False positive events, such as the event in base 44 that arose from a single anomalous replicate, are removed. Similarly, qualitatively small true positives are also removed: the downswing at length 88 nt in base 14 and the downswings around 100 nt in bases 28-29. We previously observed that base 14’s downswing is likely non-spurious and marks a new discovery. Similarly, the downswings in bases 28-29 are attributed to their pairing off before the final structure. These removals underscore the tradeoff that while higher thresholds lower false positives, they also turn true positives into false negatives. We chose a large increase but retained many of the originally detected events, suggesting that large events have a wide acceptable range of threshold values.

Conversely, the lenient scenario (Supplementary Figure S4, left; 50% decrease in each threshold) leads to more true and false positives. The upswings in bases 26 and 33 around length 90 nt become detected. By inspection, these events seem non-spurious and occur concurrently with other similar events (Supplementary File 1) leading to the conclusion that these are previously undetected true positive events. Conversely, the lenient scenario creates potential false positive events. For example, the upswings in base 40 at 118 nt and all upswings in base 44 do not resemble upswings. The upswings in base 44 are especially misleading; one replicate has increased reactivity while the others remain flat (Supplementary Figure S4). We conclude that the lenient scenario reveals spurious events.

We chose drastic perturbations to threshold parameter values to interrogate their effect on detection rates. Many originally detected events remained in the stringent scenario and relatively few spurious events arose in the lenient scenario, suggesting that our methodology yields concordant results across a wide acceptable range of thresholds. We provide additional sensitivity analysis on window length and linear ramp thresholds in Supplementary Materials.

## 4 Conclusion

DUETT emulates human visual inspection of cotranscriptional SHAPE-Seq data in an automated, efficient and systematic manner to reduce potential user bias discover novel events that are difficult to manually detect. Cotranscriptional SHAPE-Seq complements aptamer structure and dynamics information (known from crystallography and NMR) with the ability to probe nascent RNA structures during transcription. When interpreting cotranscriptional SHAPE-Seq data, it is also important to keep in mind that halted nascent RNA structures are probed and fleeting structural changes are difficult to detect. As we demonstrate, DUETT detects many of the previously identified signatures of nascent structures within three model systems and identifies several new events absent from manual visual analysis. Experimentalists can now quickly establish transcription lengths and nucleotides of interest from reactivities to be further interpreted and developed into a structural model. Additionally, the automated analysis allows experimentalists to use their reactivity measure of choice. In this study, we chose ρ reactivities for the ability to compare reactivities between different length transcripts and across different experiments. We note that correlations within a ρ reactivity vector make interpretation of detected events harder, especially when concurrent up and down events occur, as it could be reflective of either structural changes or reactivity calculation. We hope this method is readily adopted when studying new RNA systems, or interrogating publicly available datasets in the RNA Mapping DataBase (RMDB) (Cordero *et al.*, 2012). DUETT is a powerful method to identify structural events that evade manual identification in cotranscriptional SHAPE-Seq data.

## Supplementary Materials

This supplementary file contains the methods for DUETT, results from the automated threshold selection process, results from averaging replicates then conducting event detection, additional sensitivity analysis, a table of assumptions, and a table of user-defined threshold parameters used in the SHAPE-Seq event detector manuscript. This word document also contains descriptions for Supplementary Files 1-3.

## DUETT methods

### Swing event detection is motivated by PI control

Feedback control is widely deployed throughout the process systems industries to mitigate fluctuations of key process variables about a desired system state (Minorsky, 1922). One common application is to maintain steady-state response by taking corrective action based on the measured deviation of controlled system variables from their nominal steady-state values. PI controllers scale the strength of their corrective action based on both the *proportion (P)* to, and *integral (I)* of, the measured deviation from steady state. The *P* and *I* terms provide a convenient framework for quantifying the magnitude and duration of the system’s deviation from steady state, which we use to detect significant SHAPE reactivity events.

Swing events are characterized by abrupt changes in SHAPE reactivity during transcript elongation. These events may be represented as *deviations* (△) about a constant reactivity value:

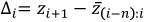

where *i* indexes transcript length, *z* is SHAPE reactivity, and 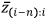 is mean reactivity within a sliding window of length n starting at position *i* − *n* and ending at position *i*. DUETT quantifies the absolute magnitude and duration of these deviations at each transcript length by adopting both *p* and *I* terms:

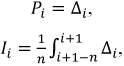

where *I* is normalized by the window size. A third quantity captures the relative magnitude (*R*) of deviations:

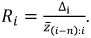

DUETT detects an upswing when each of the *P*, *I*, and *R* (PIR) values exceed set thresholds. Higher *P* and *R* thresholds require that reactivity changes are large and distinct enough from the local steady state (Supplementary Figure S1). Higher *I* thresholds ensure that deviations reflect sustained reactivity changes rather than brief noise-driven fluctuations (Supplementary Figure S1). All threshold values are positive; a zero-value denotes constant reactivity.

Downswing events are detected similarly using the additive inverse of the *PIR* thresholds. The downswing *R* threshold requires a slight modification to retain an equivalent magnitude to its upswing counterpart:

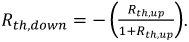

For example, an upswing event with 50% increase relative to the local steady-state reactivity (*R_th,up_* = 0.5) is of equivalent magnitude to a downswing event with 33% decrease (*R_th,down_* = 0.33).

We introduce two additional threshold parameters to further mitigate the impact of minor fluctuations. We remove swing events that are shorter than a specified *duration threshold* as we assume that structurally informative events generally persist for longer durations (Supplementary Table S1). Conversely, we merge swing events separated by a distance less than a *gap threshold* to reject noise-driven fluctuations that intersperse real swing events.

### Automated parameter selection for swing event detection

DUETT provides a method to automatically select *PIR* thresholds for any given dataset. Due to the subjective nature of event detection, the automated strategy relies on a heuristic similar to the elbow rule in cluster analysis (Ketchen and Shook, 1996). The heuristic identifies a threshold combination that balances lenient with stringent thresholds. The automated search starts with low PIR thresholds where both noise and real events are detected. DUETT scans over combinations of increasing *PIR* threshold values (within a user-defined range) and records the number of detected events. We expect a sharp decrease in the number of detected events as threshold values increase, followed by a leveling off, forming an elbow. DUETT identifies the vertex of the elbow—representing detection of true events—by finding the *PIR* threshold values that correspond to a detection output closest to the origin. If needed, the automatically identified thresholds serve as a starting point for manual tuning.

### Ramp event detection using linear regression

Ramp events correspond to gradual changes in reactivity over broad stretches of sequential transcript lengths. The swing event detection method overlooks ramp events because it emphasizes rapid changes in reactivity over few transcript lengths. Instead, DUETT detects ramp events using a strategy based on linear regression. Given a user-specified ramp length corresponding to the minimum expected ramp length, lines of the form *y* = *βx* are fit within windows sliding down each column of the SHAPE-Seq data matrix. Here, *x* is a vector of sequential integers with length equal to the *ramp length, y* = *βx* is the measured reactivities within the corresponding window, and *β*; is a regression coefficient.

A ramp is detected when a fitted line passes three user-specified thresholds: a maximum *p-value* calculated from a one sample t-test for the *regression coefficient*, a minimum regression coefficient (*β*), and a minimum *Durbin-Watson Statistic (DWS)*. Manual tuning of these thresholds is required to confidently detect ramp events. Fortunately, these parameters are readily interpretable (Supplementary Figure S1). The *p*-value threshold controls the robustness of event detection against measurement noise; low values improve specificity at the expense of sensitivity. The *β* threshold constrains ramp steepness; high values exclude relatively flat ramp events with low average change in reactivity. Finally, the *DWS* threshold tunes the selectivity of ramp versus swing event detection. Swing events yield strong positive autocorrelation among residuals about the regression line (Supplementary Figure S1, right panel of row 4), while residuals associated with true ramp events are uncorrelated. A *DWS* threshold of unity precludes misclassification of swing events as ramps by filtering events whose sequential residuals are positively correlated (Nerlove and Wallis, 1966).

### Identifying concurrent events

Multiple events across different nucleotide positions detected at similar transcript lengths likely reflect a common structural change. For example, a pair of upswings independently detected at two different nucleotide positions are involved in the same structural rearrangement if they occur at approximately the same transcript length. We label such instances as concurrent events by applying a *concurrency distance* proximity threshold.

### Computational development and graphical user interface

DUETT was programmed in the freely available statistical software environment R and RStudio. We provide the source code and a graphical user interface as an R Shiny app located at https://github.com/BagheriLab/DUETT. The app facilitates parameter tuning by continually updating figures as parameter values are varied by the user. The app also provides a suite of formatting tools for generating appropriate figures and tables.

### Additional sensitivity analysis Window length

We explore how window lengths affect event detection. Generally, longer window lengths lead to higher sensitivity (more true and false positives) because reactivities from shorter/earlier transcript lengths affect the average window used in *PIR* equations. With a 100% longer window (Supplementary Figure S5 right), this effect is shown with base 14, resulting in detected upswings. As a tradeoff, spurious events are included such as the downswing in base 37 (Supplementary Figure S5 right). On the other hand, a shorter window generally decreases sensitivity (Supplementary Figure S5 left). The upswings in base 40 after 75 nt are no longer steep enough to be included and the overall number of swing events has decreased.

A longer window does not globally increase event detection sensitivity as evident in false negatives for downswings in base 14 at length 87 nt (Supplementary Figure S5). This observation is due to two effects. First, the transcript lengths before 87 nt have an upward trend, and the longer window length averages over lower reactivity transcript lengths and causes the downswing to appear less significant. Second, the integral length for *I* defaults to the same value as the window length, and longer integral lengths lower sensitivity. Instead, specifying a shorter window length (5) for the *I* length generally increases sensitivity and recapitulates the downswings in bases 11 and 14 (Supplementary Figure S6).

This analysis demonstrates that longer windows and shorter windows generally enhance and decrease sensitivity, respectively, but scenarios exist that go against this rule. Our detailed analysis serves as a cautionary tale that there are tradeoffs between identifying false and true events.

### Durbin-Watson Statistic

We provide sensitivity analysis for the Durbin-Watson statistic (*DWS*). We recommend a default setting between 1 and 1.25 as lower values traditionally correspond to a positive correlation, and we test a lenient (0.1) and a stringent threshold (1.5). As expected, the lenient and stringent thresholds have more and fewer detected ramps, respectively (Supplementary Figure S7). The lenient threshold allows linear ramps where the residuals are not uniformly distributed down the length of the ramp. For example, the downramp in base 14 has points clustered above and below the ramp, and we argue that this pattern does not conform to our expectations of a ramp and appears more like a downswing (it is detected as a downswing). In contrast, the upramp in base 41 appears genuine but is removed with the stringent *DWS*. Though minor, the flat region around length 50 nt creates a sequence of negative residuals that correspond to a positive autocorrelation, which fails the stringent *DWS*. However, we observe that the upramp in base 42 is preserved because it lacks a flat region as large as in base 41. These three examples showcase how the *DWS* threshold parameter removes ramps that resemble swing events

Overall, we apply large threshold changes and most qualitatively large events are still identified in all scenarios. This demonstrates that large and clean events can be insensitive to threshold changes, and the true positive-false positive tradeoff is applicable to qualitatively small or noisy events. Consequently, it is likely that the SHAPE-Seq event detector identifies large RNA structural events with similar accuracy as a human, but the detector can also systematically identify small events whereas a human might not. Similarly, RNA structural events are not clearly defined and encompass a degree of subjectivity meaning that tradeoffs between identifying true positives and avoiding false positives/negatives must be considered. Altogether, these complexities justify the need for our quantitative and systematic approach.

## Supplementary File descriptions

Supplementary Files 1-3 contain the profiles for each nucleotide in *E. coli* SRP and the *B. cereus* crcB fluoride riboswitch datasets without fluoride and with fluoride, respectively. Upswings, downswings, upramps, and downramps are shown with red diamonds, blue diamonds, red lines, and blue lines, respectively. Concurrent events are denoted with a dotted green line.

---

**Figure S1.**
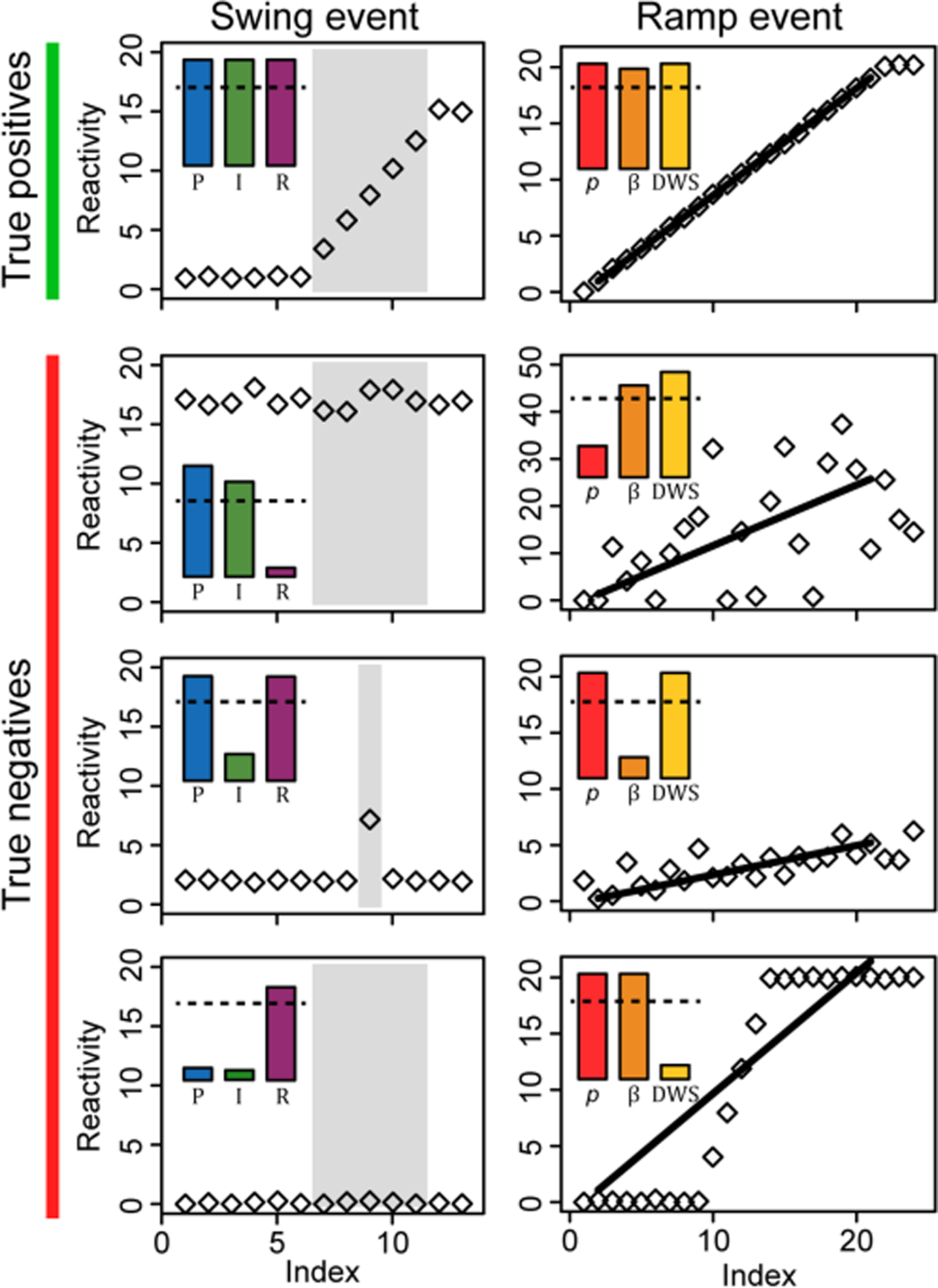
Three threshold parameters detect true positive events from true negative ones. Swing and ramp events differ in terms of event length and have their own set of threshold parameters for detection. DUETT detects swing events using a strategy inspired by control theory where proportional (*P*) and relative (*R*) thresholds filter out noise at low and high magnitudes, respectively, and the integral (*I*) threshold filters out anomalous high values that persist for short durations. In silico illustrations and representative PIR values are shown for events in the region of interest (grey). DUETT detects ramp events using linear regression, where the ramp p-value (*p*), ramp slope (*β*) and Durbin-Watson statistic (*DWS*) thresholds filter out high noise, low slope ramps, and swing events that may be erroneously classified as ramps, respectively. We show p, β and values for the fitted line (solid line). p is shown as ‐log_10_(p) so larger values pass the set threshold values (dotted line).

**Figure S2.**
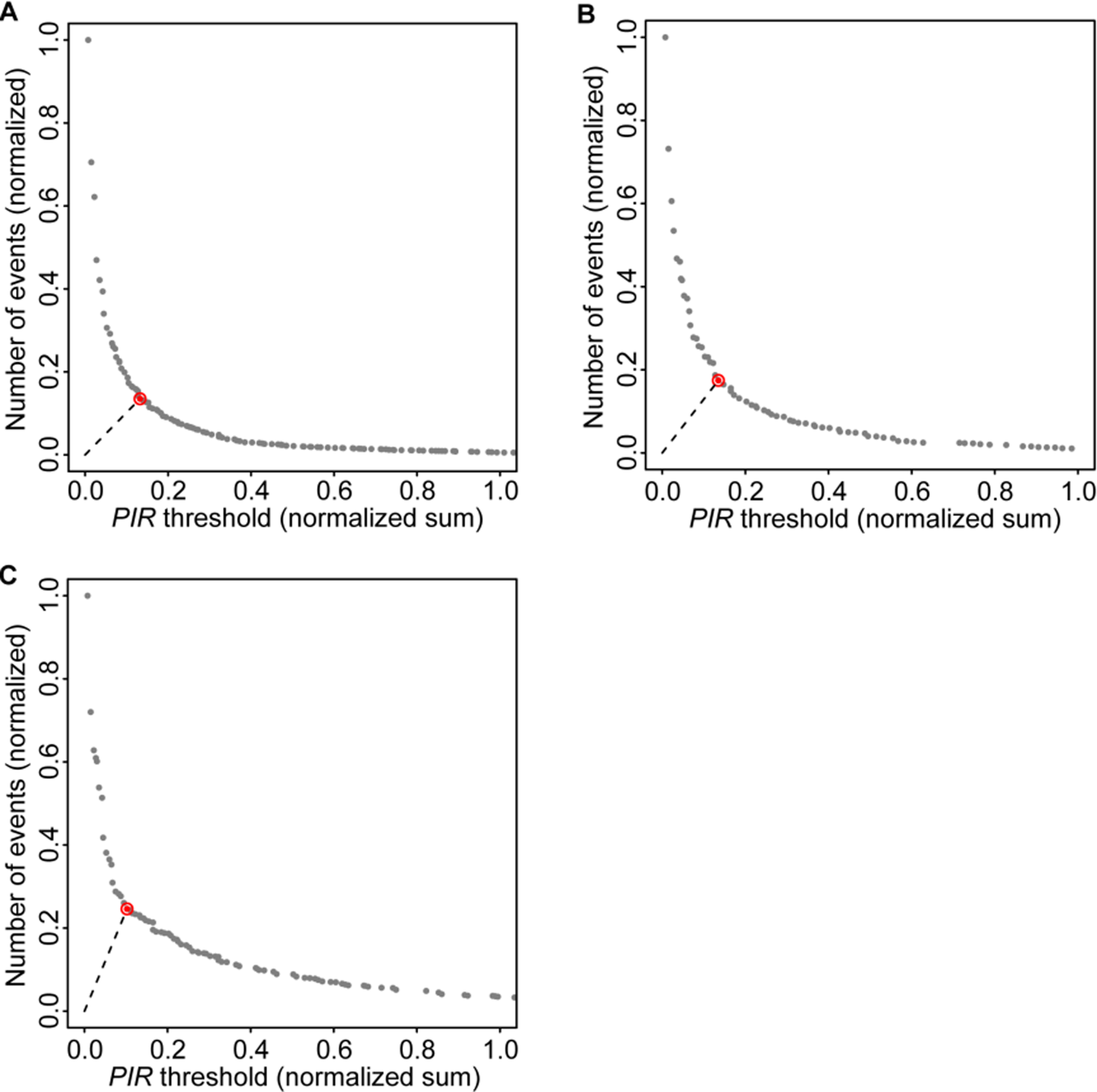
Automated PIR threshold selection identifies the balance between too lenient and too stringent parameter sets. DUETT provides an automated method to select appropriate *PIR* thresholds for A) the SRP RNA, B) the fluoride riboswitch under 0mM NaF, and C) the fluoride riboswitch under 10 mM NaF datasets. After scanning over potential combinations of *PIR* threshold values, DUETT identifies the threshold set closest to the origin (red point with dotted line from origin). This point corresponds to the elbow criterion identifying the point when spurious detection is avoided while retaining true events. The horizontal axis is the sum of all three *PIR* thresholds and both axes are normalized. This calculation is done in 4-dimensions (*P*, *I*, *R*, and number of events) and the selected threshold values may not appear closest to the origin when plotted in 2D.

**Figure S3.**
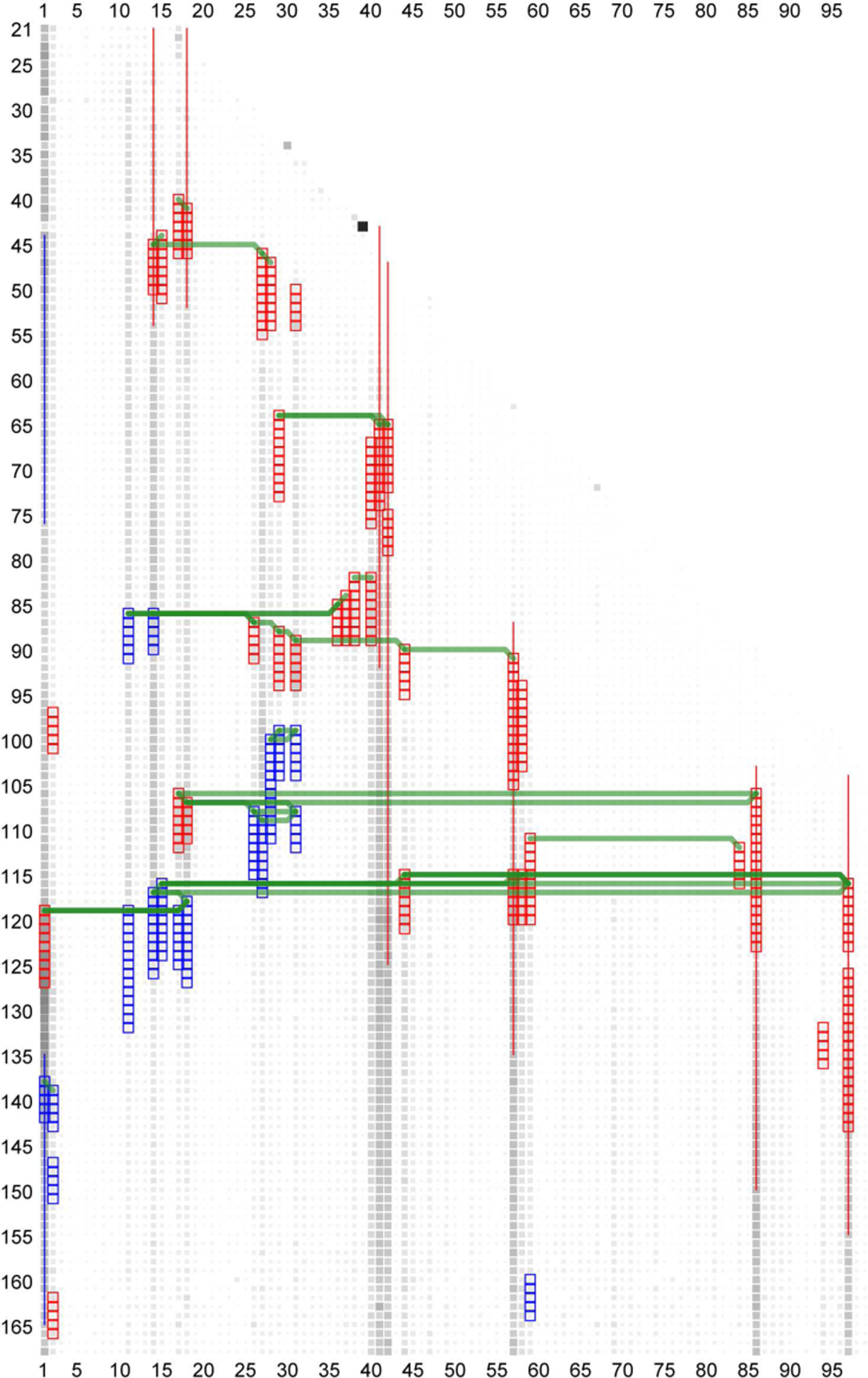
Similar results are created when applying DUETT to the average of replicates. Figure 1 shows results from identified events that are shared in each of the three replicate datasets on the SRP RNA. Here, all three replicates are averaged then event detection is conducted. The *PIR* thresholds are slightly more stringent than in Figure 1 because few events pass all thresholds and are conserved in all replicates. Each *PIR* threshold was increased by 0.1.

**Figure S4.**
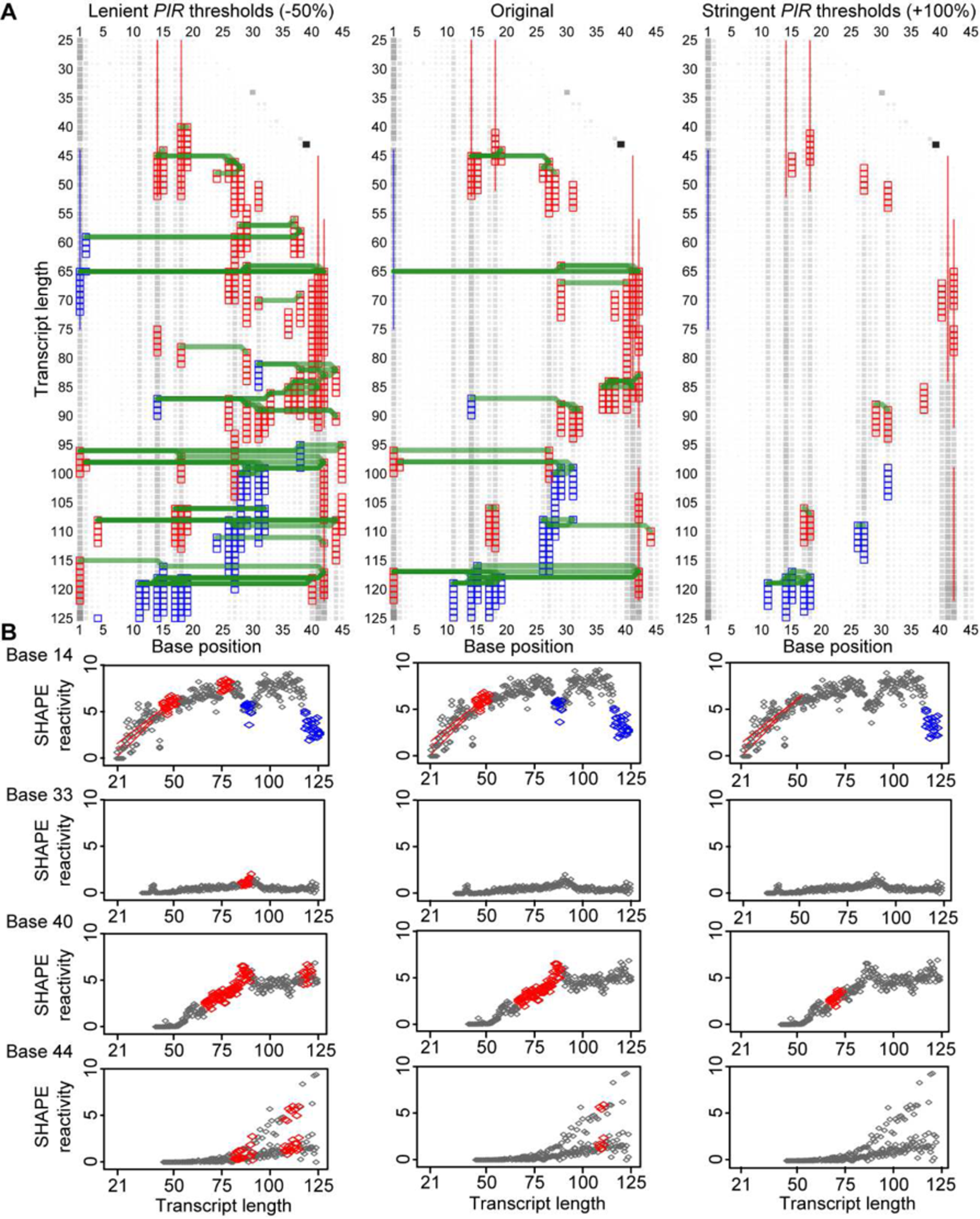
Sensitivity analysis of user-defined threshold values illustrates the tradeoff between true positive and false positive/negative detection. A) The original *PIR* thresholds (center) are compared to a lenient scenario (left, 50% decrease in all threshold values) and a stringent scenario (right, 100% increase in all threshold values) on the SRP dataset. In general, lenient thresholds increase sensitivity towards small magnitude events, but false positive events are detected with greater likelihood. Conversely, stringent thresholds have fewer false positive events, but the likelihood of detecting false negative events increases. Large magnitude events tend to be insensitive to large threshold deviations. B) Individual nucleotide positions are highlighted where reactivity profiles of all three replicate datasets are shown. Events are annotated as in Figure 1 except that reactivities and swing events are shown with diamonds.

**Figure S5.**
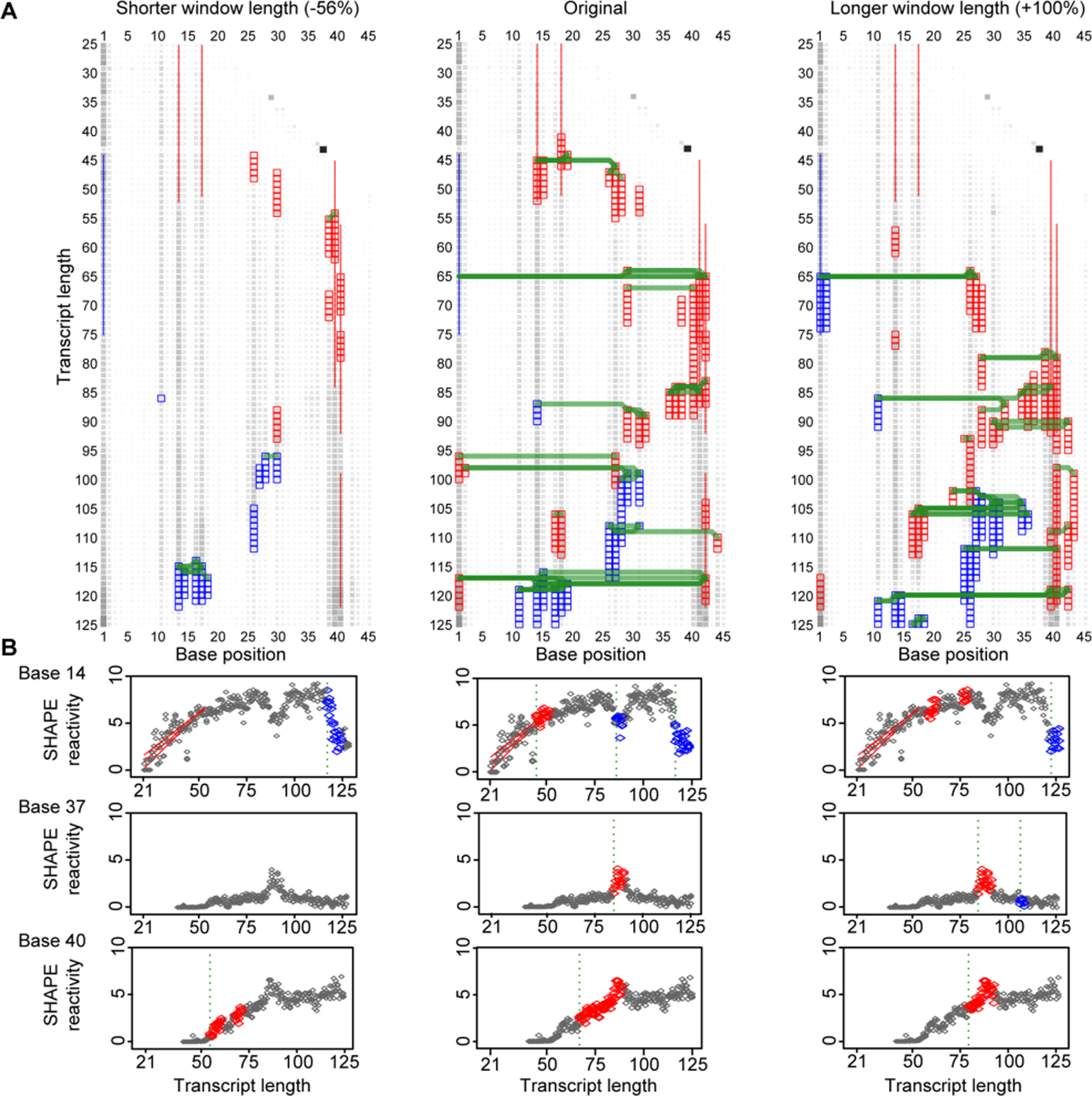
Sensitivity analysis of window length. Window length effect is shown A) in general and B) with specific examples on the SRP dataset. The window length determines the number of positions that are averaged together before *PIR* calculation. Longer lengths generally lead to higher sensitivity of true positives because the longer window retains earlier reactivities. In contrast, shorter lengths are less sensitive and generally lead to fewer events and false positives/negatives.

**Figure S6.**
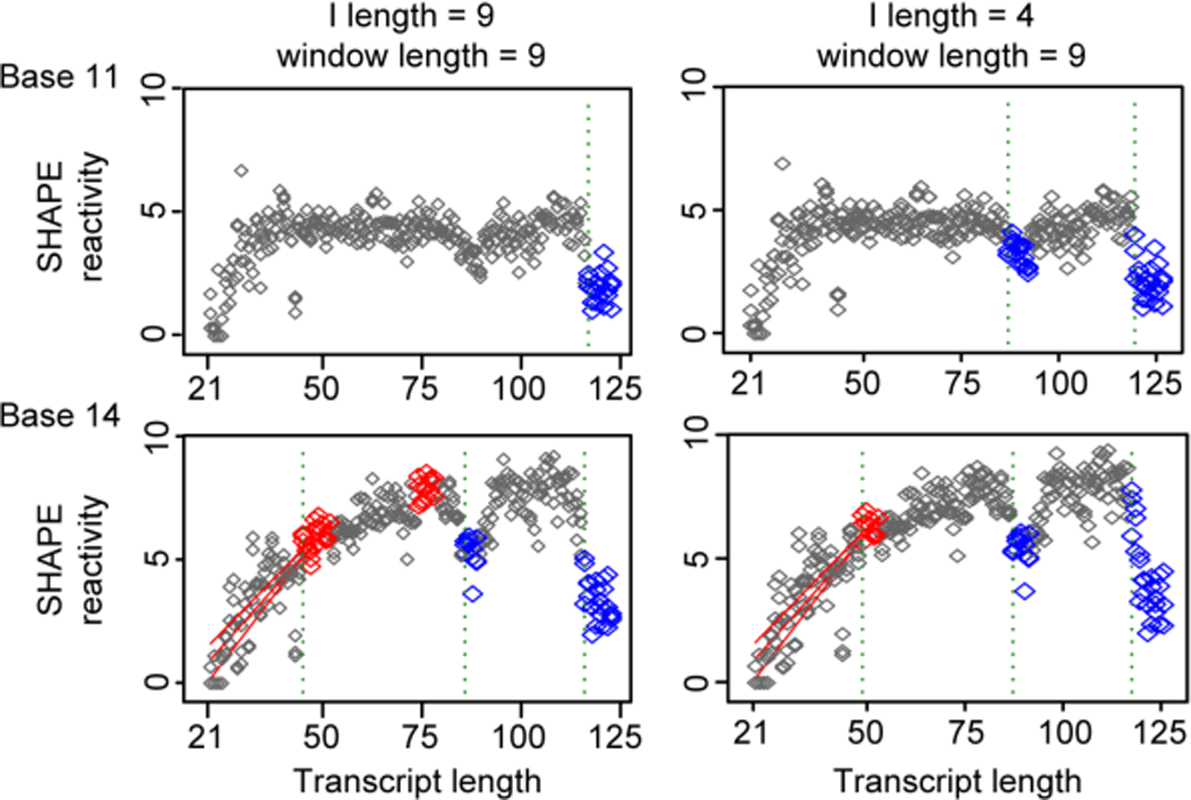
Higher *I* length generally lowers sensitivity and longer window length does not always raise sensitivity. By default, *I* length is the same as the window length (left) but can be specified separately (right). Lower and higher values of *I* length generally increase and lower sensitivity, respectively. Examples are drawn from the SRP dataset.

**Figure S7.**
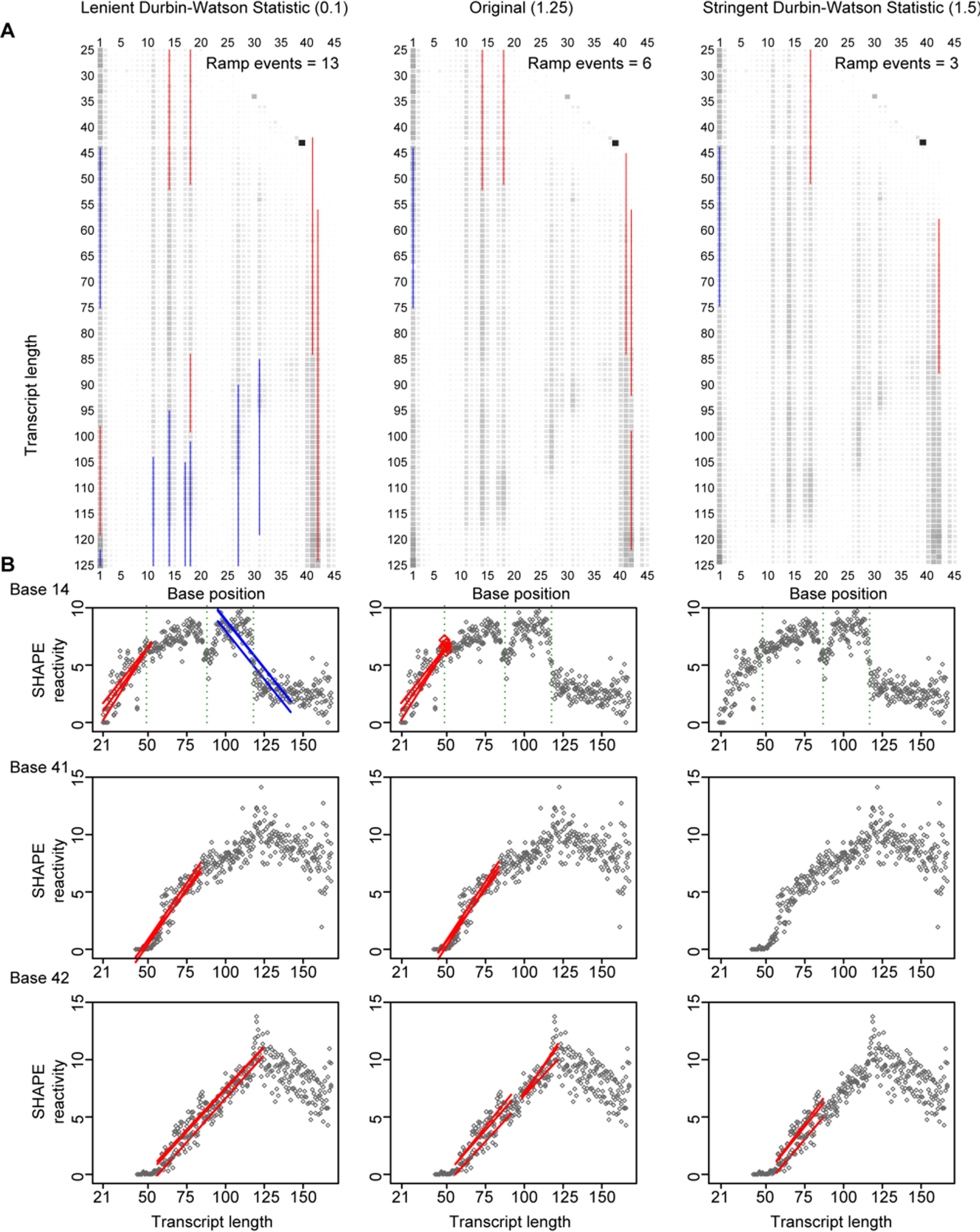
Sensitivity analysis of the Durbin-Watson statistic. *DWS* ranges from 0-4 where 2 represents our ideal scenario of no autocorrelation in the residuals. We present effects of lower and higher *DWS* thresholds and compare the quality of fitted linear ramps on the SRP dataset. Generally, lower *DWS* thresholds are more lenient where lines are fitted on less line-like segments. To simplify visual analysis, all swing events were removed.

**Table S1.**
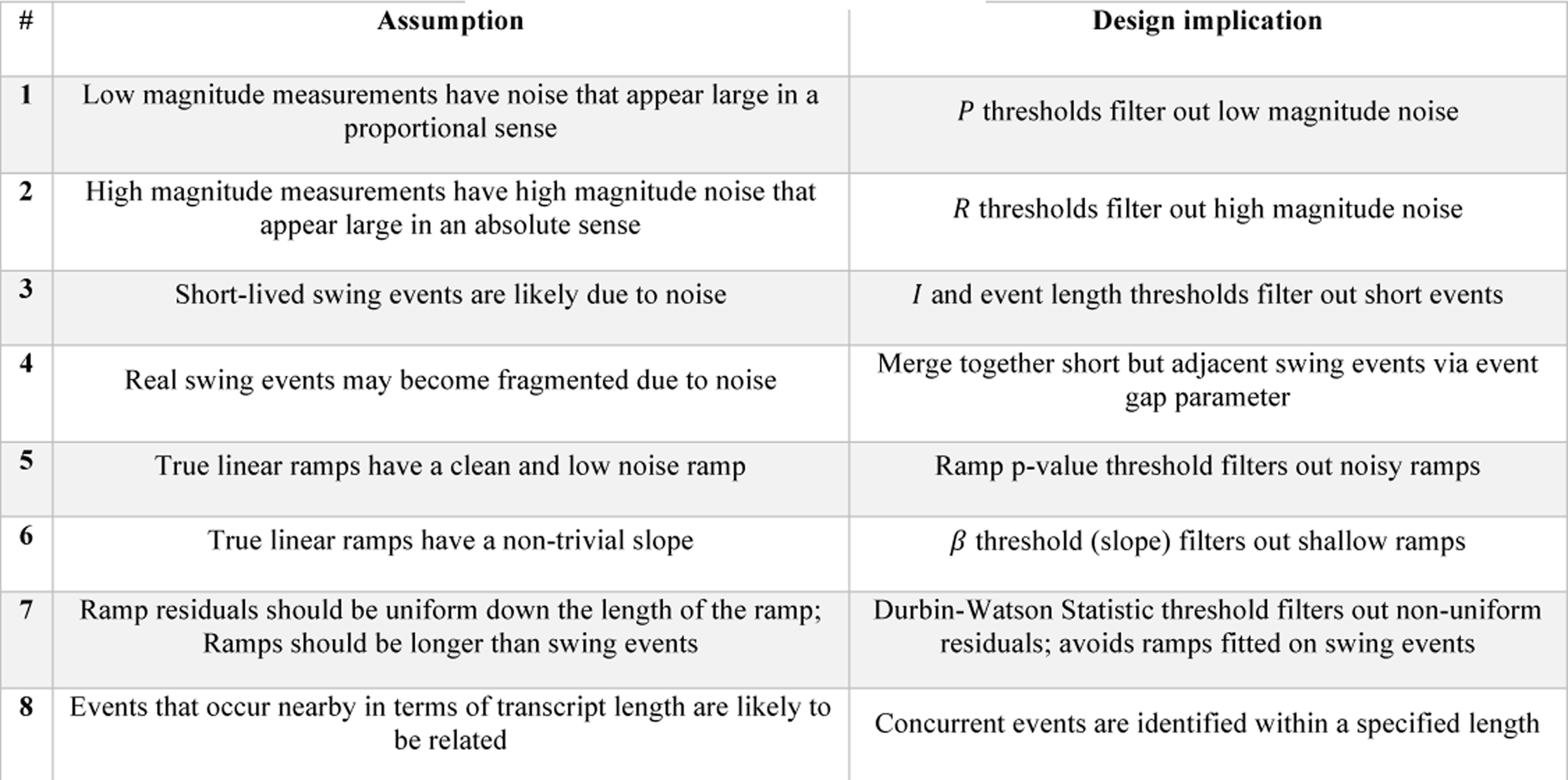
Explicit assumptions and design implications in DUETT

| # | Assumption | Design implication |
| --- | --- | --- |
| 1 | Low magnitude measurements have noise that appear large in a proportional sense | $P$ thresholds filter out low magnitude noise |
| 2 | High magnitude measurements have high magnitude noise that appear large in an absolute sense | $R$ thresholds filter out high magnitude noise |
| 3 | Short-lived swing events are likely due to noise | $I$ and event length thresholds filter out short events |
| 4 | Real swing events may become fragmented due to noise | Merge together short but adjacent swing events via event gap parameter |
| 5 | True linear ramps have a clean and low noise ramp | Ramp p-value threshold filters out noisy ramps |
| 6 | True linear ramps have a non-trivial slope | $\beta$ threshold (slope) filters out shallow ramps |
| 7 | Ramp residuals should be uniform down the length of the ramp;<br>Ramps should be longer than swing events | Durbin-Watson Statistic threshold filters out non-uniform residuals; avoids ramps fitted on swing events |
| 8 | Events that occur nearby in terms of transcript length are likely to be related | Concurrent events are identified within a specified length |

**Table S2.** Automated *PIR* and user-defined linear ramp threshold parameters for the SRP and fluoride riboswitch examples

| *PIR* |  |
| --- | --- |
| Window length | 9 |
| $P$ | 0.3 |
| $I$ | 0.025 |
| $R$ | 0.1 |
| $I$ length | default |
| Noise length | 4 |
| Event gap | 1 |

| Ramp |  |
| --- | --- |
| Ramp length | 30 |
| $p$-value | 1.00E-04 |
| $\beta$ | 0.15 |
| $DWS$ | 1.25 |
| Concurrent distance | 1 |

F riboswitch settings (0 mM NaF left, 10 mM NaF right)
| *PIR* |  |  |
| --- | --- | --- |
| Window length | 9 | 9 |
| $P$ | 0.35 | 0.3 |
| $I$ | 0.05 | 0.05 |
| $R$ | 0.025 | 0.025 |
| $I$ length | default | default |
| Noise length | 4 | 4 |
| Event gap | 1 | 1 |

| Ramp |  |  |
| --- | --- | --- |
| Ramp length | 30 | 30 |
| $p$-value | 1.00E-04 | 1.00E-04 |
| $\beta$ | 0.15 | 0.15 |
| $DWS$ | 1.25 | 1.25 |
| Concurrent distance | 1 | 1 |

